# CellScape: Protein structure visualization with vector graphics cartoons

**DOI:** 10.1101/2022.06.14.495869

**Authors:** Jordi Silvestre-Ryan, Daniel A. Fletcher, Ian Holmes

**Affiliations:** Department of Bioengineering, University of California, Berkeley, 94720, USA

## Abstract

**Motivation:** Illustrative renderings of proteins are useful aids for scientific communication and education. Nevertheless, few software packages exist to automate the generation of these visualizations.

**Results:** We introduce CellScape, a tool designed to generate 2D molecular cartoons from atomic coordinates and combine them into larger cellular scenes. These illustrations can outline protein regions in different levels of detail. Unlike most molecular visualization tools which use raster image formats, these illustrations are represented as vector graphics, making them easily editable and composable with other graphics.

**Availability and Implementation:** CellScape is implemented in Python 3 and freely available at https://github.com/jordisr/cellscape. It can be run as a command-line tool or interactively in a Jupyter notebook.

## Introduction

Molecular visualization plays a fundamental role in structural biology, and generally involves a balance between realistic depictions of atomic details and abstractions designed to highlight structural features. For example, proteins are often shown as ribbons of secondary structure with most or all sidechains hidden. One class of representations are “non-photorealistic renderings", which abstract some molecular detail to convey a structure’s general shape and form (Goodsell *et al*., 2019).

In contrast to the large number of general purpose molecular viewers, e.g. PyMOL (DeLano, 2002) or Chimera (Pettersen *et al*., 2004), there are relatively few tools designed to generate such approximate molecular illustrations. Notable exceptions include Illustrate (Goodsell *et al*., 2019), which renders individual proteins, and CellPAINT (Gardner *et al*., 2021), which is designed for dense mesoscale scenes.

Here we introduce CellScape, a flexible software tool designed to generate data-driven 2D cartoons from protein structures. CellScape can visualize structures at different levels of atomic detail, from a fine-grained view showing individual residues all the way up to a flat outline capturing only the molecule’s silhouette. Unlike most molecular visualization tools, CellScape can save images as vector graphics, allowing for easy manipulation with software such as Inkscape or Adobe Illustrator. Additionally, CellScape can combine cartoons to generate illustrative cellular landscapes, highlighting domain and geometry differences between proteins. These two usage modes for building and combining cartoons are available as command-line tools (`cellscape cartoon` and `cellscape scene`), and cartoons can also be created interactively through a Jupyter notebook interface. In this note, we describe CellScape’s key features and highlight some examples of its usage (Figure 1).

**Fig. 1.**
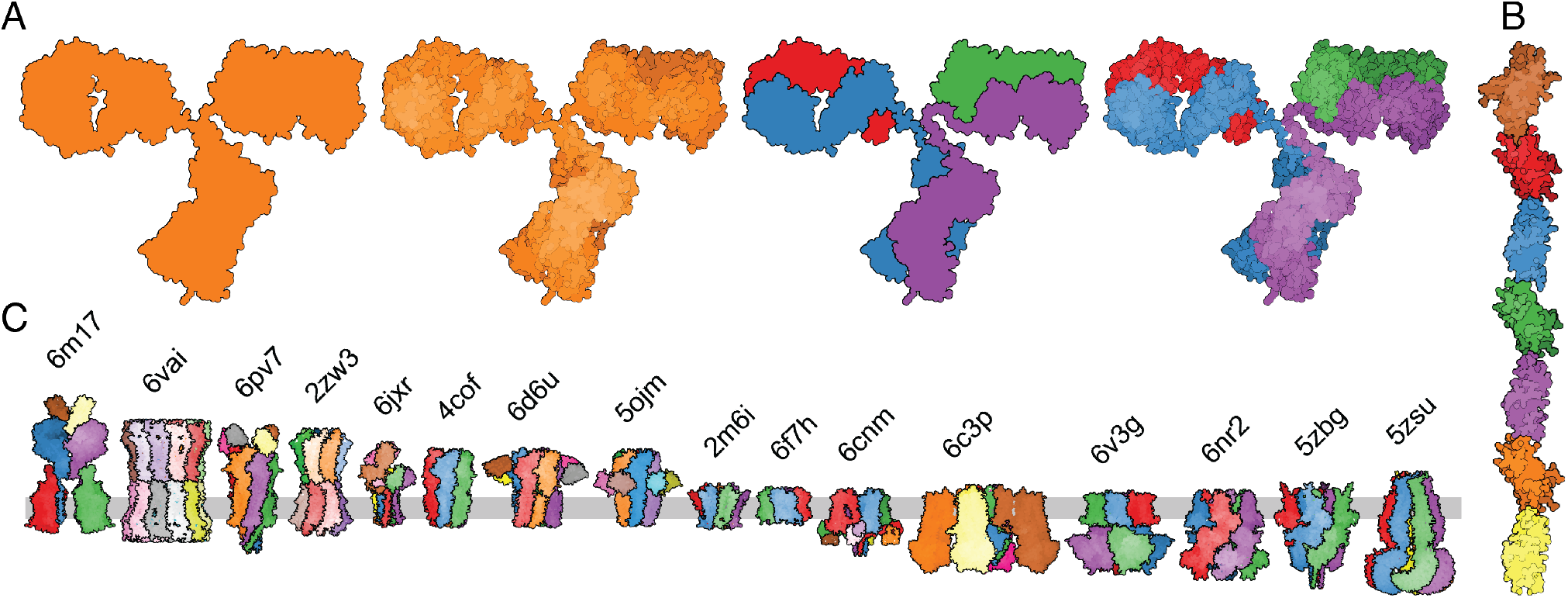
Examples of styles and renderings using CellScape for (A, B) a single protein and (C) a cellular scene. A) From left to right, illustrations depict an immunoglobulin structure (PDB 1igt) using: an outline of the entire protein, depth slices through the protein, outlines of each chain, and outlines of each residue that are colored by chain. B) Model of CEACAM5 (based on PDB 1e07) with domain bounds parsed from UniProt. C) Sample molecular scene depicting an assortment of human membrane proteins with solved structures. Placement in the membrane was determined from the Orientation of Proteins in Membranes database (Lomize et al., 2012). Labels indicate corresponding PDB ID and were added automatically with CellScape.

## Key features

### Cartoon generation

CellScape visualization requires at minimum atomic coordinates in PDB or mmCIF format, which are parsed using Biopython (Hamelryck and Manderick, 2003). If run through a Jupyter notebook, an NGLView (Nguyen *et al*., 2018) widget allows for interactive rotation of the molecule by the user. Otherwise, the protein may be oriented separately in PyMOL or Chimera, and the camera view matrix exported to a file.

Once the desired orientation is chosen, the atomic coordinates are projected down to two dimensions. In this 2D view each atom can now be represented as a disk, and each residue as the union of all its atoms. Instead of outlining each residue separately, CellScape can also outline larger groupings (e.g. one outline for each chain, as in Fig 1A). Given a UniProt accession number, CellScape will download the record in XML format and extract domain and cellular topology annotations, which can also be incorporated into the visualization (e.g. one outline for each domain, as in Fig 1B).

In building these flat 2D outlines, the depth of each atom is still tracked, to ensure that overlaps between regions are properly occluded. As an alternative to a single flat outline, the protein can also be represented using a series of contours which slice along the Z-axis (into the screen). In views such as this, which more closely represent 3D structure, depth-based edges and shading can be used to mimic 3D lighting effects often used in molecular visualization (Tarini *et al*., 2006).

### Scene generation

After cartoons have been generated from individual molecules, CellScape can compose them together into a larger visualization. The `scene` program takes a list of cartoons (in the form of Python objects exported by `cartoon`) and draws them together on the same set of axes. Relative protein scales are preserved, such that molecule sizes can be accurately compared. This can be used simply to lay out molecules next to each other (Figure 1A), or to depict a cellular membrane scene (Figure 1C).

If provided with expression data, CellScape can generate a random scene where each protein is sampled according to its expression frequency or stoichiometry. This comma-separated input file is analogous to the concept of “mesoscale recipes” used in the 3D visualization software CellPACK (Johnson *et al*., 2014). This allows CellScape to make representative visualizations of proteins on the cell surface, which we have done for several human cell types in a study of extracellular protein heights (manuscript in preparation).

## Conclusion

CellScape is a versatile tool for generating and composing simplified representations of protein structures. These visualizations can highlight different levels of structural detail by outlining individual residues, domains, or chains. CellScape can combine cartoons into scenes depicting proteins on a cellular membrane, or it can export them to vector art for easy editing and arrangement by the user. CellScape visualizations can thus be used as is, or as building blocks for more complex representations of the cellular mesoscale.

